# *Gymnosiphon fonensis* (Burmanniaceae) a new Critically Endangered species from Simandou, Republic of Guinea, W. Africa

**DOI:** 10.1101/2023.02.17.528802

**Authors:** Martin Cheek, Barthelemy Tchiengue, Aminata Thiam, Denise Molmou, Tokpa Seny Dore, Sekou Magassouba

## Abstract

A new species of *Gymnosiphon* Blume (Burmanniaceae), *G.fonensis* Cheek is formally described from the Forêt Classee de Pic de Fon, Simandou Range, Guinee-Forestière, Republic of Guinea (Guinee-Conakry) in West Africa. The new species was formerly confused with and resembles *G. bekensis* Letouzey of central Africa in the broad flat outer tepal lobes, perianth tube >10 mm long, and (sub)sessile flower. It differs e.g. in that the length of the corolla tube, (13-)14-18 mm, exceeds the corolla diameter (10-11 mm) (vs length of the corolla tube (12 mm) < the corolla diameter (12-15 mm)), the anthers inserted c. 4 mm deep in the corolla tube (vs inserted at the corolla mouth) and the rhizome lacks scale-leaves (vs scale leaves present). *Gymnosiphon fonensis* is the first known species of its genus and family in which secondary pollen presentation has been recorded. The species is known from five sites, all with threats, in a single threat-based location, accordingly it is assessed as Critically Endangered (CR B1ab(iii)) using the IUCN 2012 standard, making it the most threatened species of *Gymnosiphon* in continental Africa. The new species is illustrated by colour photos and line-drawings and is mapped. An identification key is provided to the ten species of the genus now known from Africa-Madagascar.

## INTRODUCTION

The Republic of Guinea (245,857 km^2^), also known as Guinea-Conakry, formerly Guinee Française or French Guinea is situated in West Africa. The country is dominated by the Guinea Highlands. They form or influence the vegetation types that give Guinea its diverse flora and have many narrow endemics. The Highlands are the highest and most extensive in coastal continental Africa west of the Cameroon Highlands which lie 2000 km to the east. The Guinea Highlands rise to between 1000-2000 m above sea-level and are divided into two portions. In the northwest, the Fouta Djalon highlands are entirely confined to Guinea. The geologically different Loma-Man Highlands to the southeast, in contrast, while mainly falling in Guinea, extend from Mts Loma in northern Sierra Leone to the Man Mts of western Ivory Coast. Major rivers such as the Niger, Senegal and Mano arise in the Highlands, hence the country is known as the “water tower” of West Africa. Rainfall falls mainly in May-October. The highest rainfall occurs at the coast (4 m p.a.) falling to 1 m p.a. in the north (Couch *et al*. 2019).

Botanical surveys of Guinea began in 1837 with the collections of Heudelot (which were often misleadingly labelled as Senegambia). During the French colonial period (1898-1958) sampling was expanded by collectors such as Chevalier, Jacques-Felix, Adam and Schnell who also published new records and taxa new to science based on their collections and those of others. The culmination of this work was the Flore (Angiospermes) de la Republique de Guinee (Lisowski 2009) which is founded on the Flora of West Tropical Africa (Keay & Hepper 1954-1972).

Since 2009, floristic surveys made in Guinea mainly by the National Herbarium of Guinea (HNG), together with colleagues from RBG, Kew (K) and Missouri Botanical Garden (MO) has resulted in the identification of new species for science in many different families of vascular plants: Apocynaceae (Goyder 2009), Acanthaceae (Champluvier & Darbyshire 2009; Darbyshire *et al*. 2011), Burmanniaceae (Cheek & van der Burgt 2010), Cyperaceae (Rosen 2010; Bauters *et al*. 2018; Verloove *et al*. 2019), Oleaceae (Jongkind 2010), Convolvulaceae (Breteler 2010; Breterler & Wieringa 2018), Leguminosae (van der Burgt *et al*. 2012; van der Burgt *et al*. 2018), Orobanchaceae (Fischer *et al*. 2012), Malvaceae/Sterculiaceae (Jongkind 2013), Lecythidaceae (Prance & Jongkind 2015), Eriocaulaceae (Phillips & Mesterházy 2015; Phillips *et al*. 2018); Dennstaedtiaceae (Jongkind & de Winter 2015), Peridiscaceae (Breteler *et al*. 2015), Euphorbiaceae (Cheek *et al*. 2016); Podostemaceae (Cheek & Haba 2016a; Cheek *et al*. 2017; Cheek & Lebbie 2018; Cheek *et al*. 2019a; Cheek *et al*. 2022a), Costaceae (Maas-van de Kamer *et al*. 2016); Orchidaceae (Bidault *et al*. 2016; Dubuisson & Stevart 2018), Rubiaceae (Cheek & Williams 2016; Jongkind 2018; Cheek *et al*. 2018a; Cheek *et al*. 2018b), Sapindaceae (Cheek & Haba 2016b), Aspleniaceae (Xu *et al*. 2019), Ternstroemiaceae (Cheek *et al*. 2019b), Lamiaceae (Phillipson *et al*. 2019); Rutaceae (Cheek *et al*. 2019c), Iridaceae (van der Burgt *et al*. 2019); Malvaceae s.s. (Cheek *et al*. 2020a); Poaceae (Xanthos *et al*. 2020; Xanthos *et al*. 2021); Olacaceae-Aptandraceae (Cheek *et al*. 2022b) and Melastomataceae (van der Burgt *et al*. 2022)

In total, 47 new species have been described since 2009 in the Republic of Guinea including 18 species and two genera which are strictly endemic. The publication of these new taxa shows that field inventory is still necessary for Guinea: new species are still frequently being discovered. Considering that 96% of the original forest vegetation (a major habitat for threatened plant species) was lost by 1992 (Sayer *et al*. 1992) and a further 25% of remaining forest cover was lost 2000-2018 in Guinea’s main forested region Guinee-Forestiere (Fitzgerald *et al*. 2021), it is clear that these and the other Red Listed species of Guinea are highly threatened and should be included in action plans (e.g. Couch *et al*. 2022) and conservation programmes such as Important Plant Areas (Darbyshire *et al*. 2017; Couch *et al*. 2019).

Achlorophyllous mycotrophic plants, often known as saprophytes, are remarkable for lacking all chlorophyll and being completely dependent on fungi for their survival. In continental Africa, individual species or entire genera which are achlorophyllous mycotrophs occur in the families *Orchidaceae, Gentianaceae*, and *Burmanniaceae*, while all members of *Triuridaceae, Afrothismiaceae* and *Thismiaceae* are achlorophyllous mycotrophs (Cheek & Ndam 1996; Cheek & Williams 1999; Cheek 2006; Sainge *et al*. 2010; Cheek *et al*. 2023). Additionally, in the Comores and Madagascar, an achlorophyllous genus of Iridaceae, *Geosiris* Baillon, occurs (Goldblatt & Manning 2010). New discoveries of achlorophyllous mycotrophs, even new genera to science, are still steadily being made across Africa and Madagascar (Nuraliev *et al*. 2016; Cheek *et al*. 2018c; Cheek *et al*. 2019d; Cheek *et al*. 2003; Cheek 2004a; Cheek & Traclet 2020).

The achlorophyllous mycotroph (Fig. 1) collected at Pic de Fon in the Simandou Range was readily identified as *Gymnosiphon* Blume due to the combination of three broad, spreading outer perianth lobes, each lobe itself with a lateral segment, and the abscission after anthesis of the upper part of the perianth tube (carrying with it the perianth lobes and stamens), leaving a naked tube surmounting the ovary. Burmanniaceae were last monographed by Jonker (1938). He included within the family the Thismiaeae and the genus *Afrothismia*, which are now treated as separate families, Thismiaceae (e.g. Maas-van der Kamer 1998) and Afrothismiaceae (Cheek *et al*. 2023). *Hexapterella* Urban, a monotypic S. American genus, is sister to *Gymnosiphon* in the molecular phylogeny of Merckx *et al*. (2006).

**Fig. 1.**
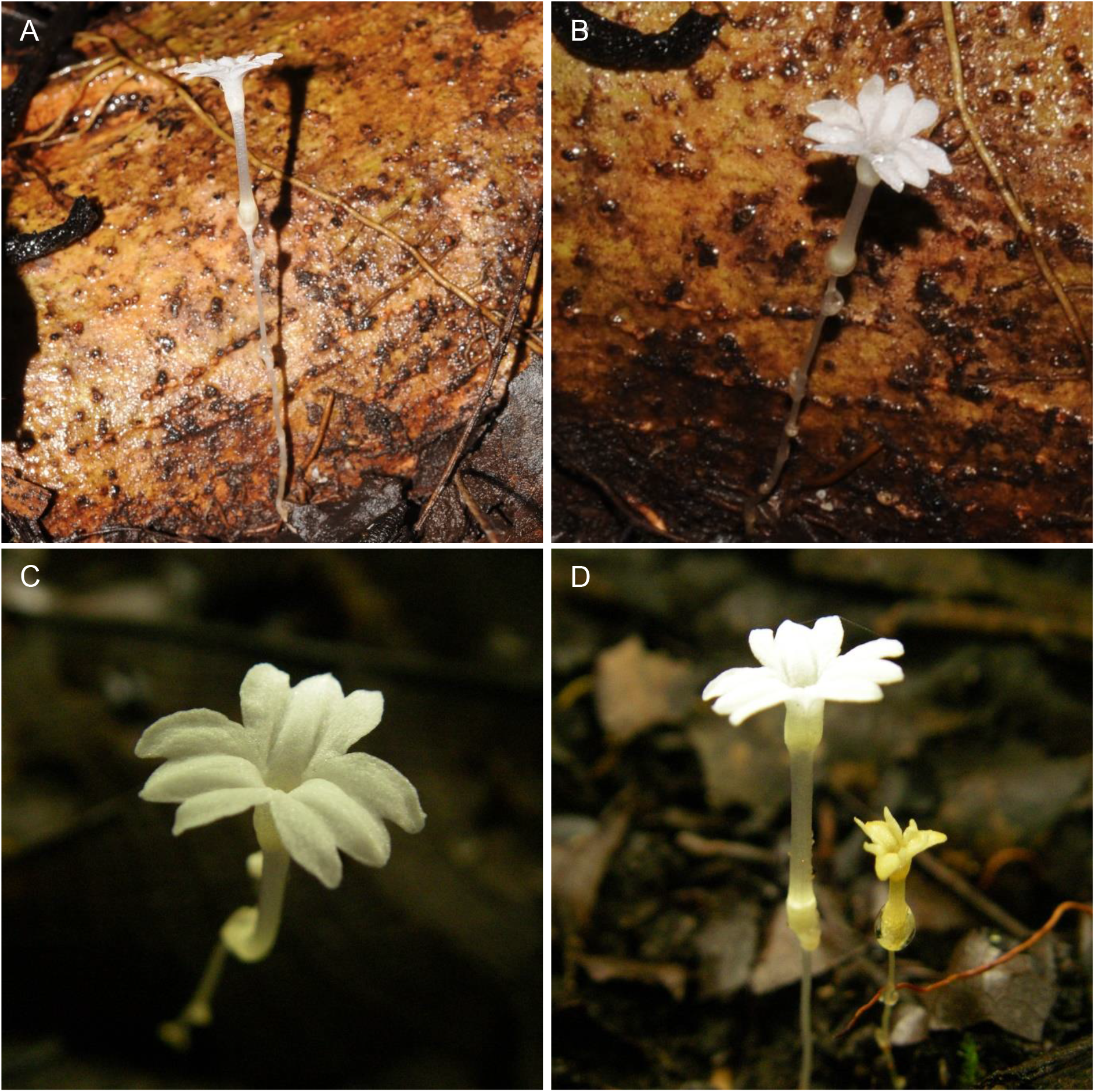
Gymnosiphon fonensis. **A.** side view of inflorescence, note cylindrical distal portion of perianth tube. **B.** flower, three-quarter view; **C.** flower showing outer perianth lobes with equal lateral and median portions; **D.** *Gymnosiphon fonensis* (left) with *Gymnosiphon samouritoureanus* (right); **A & B** photo of *Cheek* 19334 (HNG, K) by M. Cheek; **C & D** from *van der Burgt* 1274 (HNG, K) by van der Burgt

*Gymnosiphon* is pantropical with 37 accepted species (Plants of the World, continuously updated), including 17 species in the neotropics and 9 species in SE Asia, one species extending to Australia (Gray *et al*. 2019). Until the 21^st^ century, only four species were known for Africa-Madagascar, but with the discovery of *Gymnosiphon marieae* Cheek from Madagascar (Cheek *et al*. 2008), *G. afro-orientalis* Cheek from Malawi and Zambia Cheek (Cheek 2009), *G. samoritoureanus* Cheek from Guinea (Cheek & van der Burgt 2010), *G. constrictus* Maas & H.Maas from Gabon (Maas & Maas van der Kamer 2010), and *G. mayottensis* Cheek from Mayotte (Cheek & Traclet 2020), the number has more than doubled to nine.

The Pic de Fon material was initially considered to be possibly *G.bekensis* Letouzey due to the combination of a long corolla tube (> 10 mm) and the large diameter of the open flower (>10 mm), but detailed examination of material under the magnification showed it to be abundantly distinct from that species (see Table 1) and from all other species in Africa and Madagascar (see identification key below).

**Table 1.** The more significant diagnostic characters separating *Gymnosiphon fonensis* and *G. bekensis*. Data for *Gymnosiphon bekensis* was taken from Letouzey (1968) and specimens at K.

|  | *Gymnosiphon bekensis* | *Gymnosiphon fonensis* |
| --- | --- | --- |
| Geographic range | Central Africa (Democratic Republic of Congo, E. Cameroon, C, African Republic, R. of Congo) | Republic of Guinea (Forêt Classee de Pic de Fon) |
| Shape of distal portion of perianth tube | Funnel-shaped, about twice as wide at apex than the base | Subcylindrical, about as wide at apex as base. |
| Inner perianth lobes | Absent | Inserted inside perianth tube |
| Stamen insertion | At mouth of perianth tube | Inserted c. 4 mm deep inside perianth tube |
| Proportions of corolla diameter: length of perianth tube | Corolla diam. (12-15 mm) about equal or exceeding tube length (12 mm). | Corolla diam. (10-11 mm) smaller than tube length (exceeding tube length ((13-)14-18 mm). |
| Shape of outer tepal lobe apices | Acute | Rounded |
| Comparison of lateral with median portion of the outer repal lobes | Slightly longer than median, axis incurved not straight | Lateral and median equal/subequal |
| Stigma shape and posture | Funnel-shaped, erect | Obtriangular, horizontal |
| Rhizome scale-leaves | Conspicuous, forming a loose rosette around peduncle base | Absent |

The number of species described as new to science each year regularly exceeds 2000, adding to the estimated 369 000 already known (Nic Lughadha *et al*. 2016; Cheek *et al*. 2020b). Only about 7% of plant species have been assessed and included on the redlist using the IUCN (2012) standard (Bachman *et al*. 2019), but this number rises to 21–26% when additional evidence-based assessments are considered, and 30–44% of these assess the species as threatened (Bachman *et al*. 2018).

Newly discovered species such as *Gymnosiphon fonensis*, reported in this paper, are likely to be threatened, since widespread species tend to have been already discovered (Cheek *et al*. 2020b). This makes it all the more urgent to discover, document and protect such species before they become extinct as are several achlorophyllous mycotrophs in Africa such as *Oxygyne triandra* Schltr. (Cheek & Cable 2000; Onana & Cheek 2011; Cheek *et al*. 2018) and *Afrothismia pachyantha* Schltr. (Cheek *et al*. 2019d) both in Cameroon. In E Africa, *Afrothismia baerae* Cheek (Cheek 2004b) known from a single site in Kenya, has not been seen since it was published 18 years ago despite searches and may also be extinct (Luke pers. comm. to Cheek 2022). Twenty-five Guinean strictly endemic plant species are already considered possibly extinct due to habitat clearance and degradation such as by fire, mainly for small-holder agriculture and grazing (Couch *et al*. 2019), but also industrial activities such as hydro-electric dams e.g. *Inversodocraea pygmaea* G.Tayl. (Cheek *et al*. 2017) and *Saxicolella deniseae* Cheek (Cheek *et al*. 2022a). The last species became extinct before it was published. Other recent cases of species apparently becoming extinct before they are discovered (Cheek *et al*. 2018d; Cheek *et al*. 2021; Moxon-Holt & Cheek 2020; Cheek & Luke 2022; Cheek *et al*. 2022c) show the importance of finding and publishing species before it is too late, so that they can be assessed for their extinction risk and conserved,

## MATERIALS & METHODS

During preparation for fieldwork in the Boyboyba forest of Simandou in late 2022, a review was made of the threatened species known to be present there from former surveys (in 2008) including *Gymnosiphon “bekensis”*. The massive apparent geographical disjunction (exceeding 2000 km) between the Guinea subpopulation and the Congolian population of *Gymnosiphon bekensis* appears to be unmatched in the genus in Africa. It was decided to test the hypothesis that the Guinea population was morphologically different, while superficially similar to, the Congolian population. In September 2022, live material in Boyboyba (Fig. 1) was compared with the protologue of *Gymnosiphon bekensis* and several differential traits were soon detected. Additional differential traits were found once freshly collected material was studied under the microscope at K (Table 1).

Names of species and authors follow IPNI (continuously updated). Nomenclature follows Turland *et al*. (2018). Specimens were collected using the patrol method (e.g. Cheek & Cable 1997). Herbarium and spirit-preserved material was examined with a Leica Wild M8 dissecting binocular microscope fitted with an eyepiece graticule measuring in units of 0.025 mm at maximum magnification. The drawing was made with the same equipment with a Leica 308700 camera lucida attachment. Herbarium codes follow Index Herbariorum (Thiers, continuously updated). Specimens were inspected from the following herbaria: B, BM, EA, HNG, K, MAO, MO, P, SRGH, YA, WAG. The format of the description follows those in other papers describing new species in *Gymnosiphon* e.g. Cheek & Traclet (2021), terminology follows Beentje & Cheek (2003). All specimens cited have been seen. The conservation assessment follows the IUCN (2012) categories and criteria.

## RESULTS

### Key to the species of *Gymnosiphon* in Africa, Comoro Islands and Madagascar

1. Stigmas with filiform extensions exserted from flower at anthesis……………2
1. Stigmas lacking filiform extensions……………5
2. Flowers zygomorphic, inflorescence capitate, Madagascar……………*G. marieae*
2. Flowers actinomorphic, inflorescences cymose. Africa and Comoro Isl……………3
3. Perianth tube 2-5 mm long……………4
3. Perianth tube 1 mm long. Gabon……………*G. constrictus*
4. Inflorescence bracts about as long as rhachis internodes, inner tepals absent; stigmas horizontal; tepals changing from white to translucent at anthesis……………*G. mayottensis*
4. Inflorescence bracts <1/4 as long as rhachis internodes, inner tepals present; stigmas pendulous; tepals white at anthesis. R. of Guinea to DRC……………*G. longistylus*
5. Rootstock with numerous bulbils, ovary ribbed, inner tepals absent. Malawi, NE Zambia, S. Tanzania……………*G. afro-orientalis*
5. Rootstocks lacking bulbils, ovary smooth, inner tepals present. Guineo-Congolian Africa or Madagascar……………6
6. Flowers sessile or subsessile, pedicel if present to 0.5 mm long……………7
6. Flowers with a distinct pedicel 1 mm long or more……………8
7. Flowers white, perianth tube 10 mm or more long……………9
7. Flowers yellow or white, perianth tube <5 mm long. R.of Guinea & Liberia……………*G. samoritoureanus*
8. Plants commonly 15-35 cm tall; base of stem completed covered in many tens of brown leaf-scales. E. Africa……………*G. usambaricus*
8. Plants always <15 cm tall; base of stem sparsely covered, with only 3-4 leaf scales. Madagascar *……………G. danguyanus*
9. Corolla diam. 12-15 mm, >/= corolla tube length 12 mm. E.Cameroon to R. of Congo…………… *G. bekensis*
9…Corolla diam. 10-11 mm, < corolla tube length (13-)14-18 mm. R. of Guinea……………*G. fonensis*

#### ***Gymnosiphon fonensis*** Cheek, sp. nov. (Figs. 1–2)

Gymnosiphon fonensis Cheek, sp. nov. *resembles* G. bekensis *in the large diameter flowers (10 mm or more diameter), long perianth tube >10 mm long, absence of long, filamentous, stigma appendages; it differs in that the diameter of the opened corolla (10-11 mm diam.) is exceeded by the length of the corolla tube (13-)14-18 mm (not opened corolla 12-15 mm diam. exceeding or equal to the length of the corolla tube (12 mm); stamens inserted at base of distal portion of perianth tube, 4 mm from the mouth (not inserted at apex of the tube); rhizome lacking scale-leaves (not with conspicuous scale-leaves).See also Table 1 and the discussion*.

##### TYPUS

###### Republic of Guinea

Guinee-Forestière, Simandou Range, Forêt Classee de Pic de Fon, Boyboyba Forest, Oueleba (also known as Oueleba) to Wataferedou path, c. 700 m, fl. 12 September 2022, *Cheek* with Molmou 19334 (holo - K barcode K000593362; isotypes HNG, P)

##### PARATYPI

###### Republic of Guinea

Simandou Range, Pic de Fon Forêt Classee, near the northern access road to Oueleba mountain, east of Oueleba 1 camp, 706 m, fl. 24 Aug. 2008, *Tchiengue 3307* (HNG); path from Oueleba 1 to village Wataferedou, north of the path, 740 m alt., fl.fr. 31 Aug. 2008, *van der Burgt* 1274 (BR, HNG, K barcode K000460554, MO, P barcode P06800898, WAG barcodes WAG0115184 and WAG0446787); Ouelaba 1 to Wataferadou path, 750 m, fl. fr. 18 Oct. 2008, *Cheek* 13779 (HNG); north of Oueleba, forest beside the road ascending to the ridge, 875 m, fl, 20 Sept. 2008, with Pierre Haba, Pepe Haba & A. Traore, Sight Record 101. Pic de Fon area, west of the pass between villages Moribadou and Lamadou, 970 m, fl. 25 Nov. 2008, with van der Burgt, Pepe Haba, Pierre Haba, A. Traore, F. Fofana, B. Diallo, Sight Record 104. Nionsomoridou, c. 4 km due W of the town, about 100 m due S of the N2 road in gallery forest, 781 m, fl. 17 Sept. 2022., *Thiam* Sight Record (photos); Est de Pic de Fon, sud-est de Whisky 1, 864 m, fl. 18 Nov. 2008, *Traor**e,*** A. (= A.S. Goman) 161 (HNG).

##### DISTRIBUTION

This species is known from the southern half of the Simandou range in the Republic of Guinea, where it is known from five stations.

##### HABITAT

primary tropical submontane forest 700-970 m altitude.

##### PHENOLOGY

Flowering was observed from late August to November.

##### ETYMOLOGY

the epithet refers to the Forêt Classee de Pic de Fon, comprising the southern half of the Simandou range in the Republic of Guinea, from which the species was discovered and to which the species is strictly endemic on current evidence.

###### DESCRIPTION

Erect, probably perennial herb 6.5-12(−15) cm tall above roots, above ground part 5-8.3 cm tall, glabrous, achlorophyllous, white when alive. Rhizome pale yellow, vertical, cylindrical-ellipsoid, 2.5-3 x 1.25-1.5 mm (Fig. 2A-B). Base of stem lacking scale-leaves. Roots 10-17 per rhizome, vermiform, to 3.5 cm long c. 0.2 mm diam., sparingly branched (Fig. 2A-B). Tuber(s) absent. Stem (peduncle) terete, erect, single (sometimes the base of the previous season’s peduncle evident adjacent to that of current season’s (Fig. 2B).), unbranched or once branched from near the base, c. 0.75 mm in diameter, internodes (2.2-)5-7 mm long (underground) or 8-14 mm long (above ground). Scale-leaves ovate-oblong 1.1-1.5 x 0.7-0.8 mm, apex rounded-truncate, base clasping the stem for 40-50% of its circumference, below surface adpressed to the inflorescence axis, above surface spreading by c. 45°. <u>Inflorescence</u> initially single-flowered then cymose-uniparous, 1-3-flowered, flowers developing in sequence. Bracts similar to scale-leaves. <u>Pedicels</u> 0.1-0.5 mm long. Flower white or slightly yellow in bud, white at anthesis, erect, the perianth tube translucent, sometimes curved, otherwise actinomorphic, 15-20 mm long, 10-11 mm wide at anthesis, scent not detected, if present faint (*van der Burgt* 1274; Cheek pers. obs. September 2022). Perianth tube (13-)14-18 mm long, cylindrical, 1.7-1.8 mm wide, distal portion of tube (above anther insertion) broadly cylindric, 4 x 2-3 mm. <u>Outer tepals</u> 3, transversely oblong in outline, 4-5 x 4.5-6 mm, patent, with three parallel lobes, equal in length, apices rounded; lateral lobes 1.5-2 mm wide, median 2-3 mm wide. <u>Inner tepals</u> (Fig. 2E) alternating with outer, inserted inside perianth tube at (1-)1.2-1.5 mm below the sinus of the outer lobes, narrowly oblong, erect, 0.8-0.9 x 0.2 mm, apex acute. Anthers (Fig. 2 G&I). inserted 3-4 mm below the corolla mouth, at base of the distal portion of the tube, becoming attached to the style head, each 0.5-0.6 mm long, 0.8 mm wide, the two lateral, anther cells orbicular, 0.4 mm wide, immediately adjacent to each other and not separated by the connective, connective absent or inconspicuous. <u>Style</u> cylindrical, 9-10 mm long, 3-lobed at apex, stigmatic head 1.35 mm wide, the three stigmas horizontal to slightly recurved, each obtriangular, 0.7-0.8 mm long, lacking filiform appendages (Fig. 2 G-I). <u>Ovary</u> cylindric, outer surface lacking conspicuous ribs, 2 x 1.65 mm, unilocular, with 3 protoseptal, intruded parietal placentas, each placenta with 50-60 ovules; septal nectary glands six, each longitudinally ellipsoid c.0.3 x 0.2 mm., in pairs flanking the protoseptae immediately below the junction with the perianth tube (Fig. 2J). Fruit indehiscent, lacking ribs, globose-ellipsoid, 3.7-5.5 x 3-3.5 mm, walls membranous; distal perianth tube remains subcylindrical, tapering gradually towards the apex, c. (1.4)4-5 x 1-1.2 mm; style exserted 2-4 mm from perianth tube remains. Seeds oblong, 0.25-0.4 x 0.15-0.2 mm, surface reticulate.

**Fig. 2.**
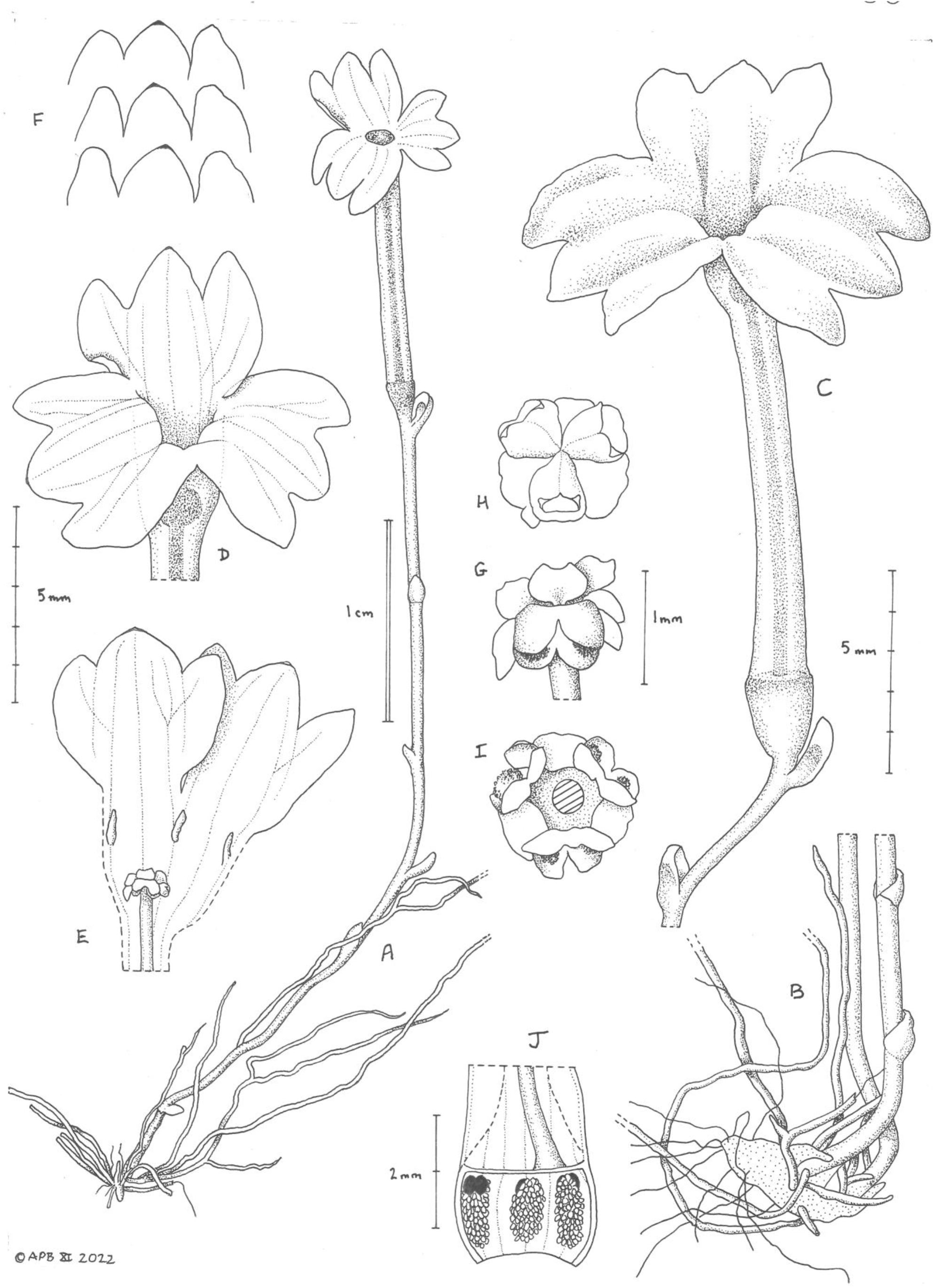
Gymnosiphon fonensis. **A**, habit, whole plant; **B**, habit, base of plant showing: rhizome (dotted) with roots and two peduncle bases; **C**, flower and rhachis; **D**, distal part of open flower; **E.** corolla tube opened (distal portion) to show inner tepals in relation to style head (with stamens attached) and stigmas; **F.** lobing of corolla lobes (from one flower); **G.** stigmatic-style head, with stigmas above, anther cells below, side view; **H.** as G but dorsal view; **I** as G but ventral view (style in transverse section). **J.** ovary opened to show the three placental masses with ovules, and (black) pairs of nectar glands. **A-I** from *Cheek* 19334 (HNG, K); **J** from *van der Burgt* 1274 (HNG, K000460554). All drawn by Andrew Brown.

###### CONSERVATION STATUS

Despite the fact that all known populations of *Gymnosiphon fonensis* sp. nov. are found in or immediately adjacent to the Forêt Classee de Pic de Fon of the southern Simandou Range, anthropogenic pressure still represents a threat and the range of the species is taken as a single threat-based location in the sense of IUCN (2012). The Pic de Fon location is set to host a major open pit iron ore mine. While the pit is not expected to directly impact sites for this species, there is a risk that despite best efforts to protect threatened species, nevertheless there will be negative impacts linked with the activities and infrastructure associated with the expected ore extraction, such as dumping of waste rock, alteration of hydrology and construction of new roads. There has already been a decline in quality of habitat e.g. due to interventions for environmental surveys such as creation of paths (Cheek pers. obs. 2021 and 2022).

Five sites are recorded based on eight specimen or photographic records, each site with 2-c.50 individuals observed. All sites are on the footprint of the project. The largest number of individuals, c.50 was recorded at the type location, a plot of c. 30 m x 10 m within the site of the Boyboyba Forest which itself measures c.0.4 km^2^ in area, while at the northernmost site, only two individuals were recorded. All five sites occur within small patches of submontane forest that appear to be dependent on streams. The forest patches are each far less than 1 km^2^ in area, however in calculating the area of occupancy (AOO) we are required to use the IUCN (2012) mandatory grid cell-size of 4 km^2^, therefore AOO for *Gymnosiphon fonensis* is calculated as 20 km^2^, below the threshold for Endangered. However, the actual area occupied by the species is likely to be far less than 1 km^2^ and is possibly less than 1 ha. For example, targeted searches at the Boyboyba Forest for achlorophyllous mycotrophs over several days in 2008 and 2022 by specialists (the authors) found *Gymnosiphon fonensis* at only a single plot of c. 30 m x 10 m (see above) while other species of achlorophyllous mycotrophs were found in multiple spots over several hectares within the forest. The sites for *Gymnosiphon fonensis* are distributed along a narrow c. 20 km north to south section of the Simandou Ridge. The extent of occurrence is calculated as 44 km^2^.

We estimate that less than 100 mature (flowering-sized) individuals are known, which would qualify the species as Endangered under Criterion D.

Since there have been past, present and future threats and AOO is above the threshold for Critically Endangered under criterion Criterion B, we here assess *Gymnosiphon fonensis* as Critically Endangered, CR B1a,b(iii). It is to be hoped that this species will be searched for and found at other locations which would allow a lower extinction risk assessment than that made here. However, since targeted searches in the correct season in 2008 outside Simandou failed to find this species, the most conspicuous species of the West African *Gymnosiphon*, it is strange that it has not been detected elsewhere before now if it indeed has a wider range. However, sites such as Mt Bero where this species might be found have not yet been thoroughly surveyed at the correct season for achlorophyllous mycotrophs.

We recommend that a management plan is developed to ensure the survival of this species and is implemented following the protocol of Couch *et al*. (2022). This should include a public sensitisation programme, annual monitoring of the *Gymnosiphon fonensis* (and also the Endangered *G. samouritoureanus*) populations to determine trends in survivorship and threats, potentially increased guarding of the forest habitat, and seedbanking. Cultivation and translocation of achlorophyllous mycotrophic flowering species such as *Gymnosiphon fonensis* has not yet been achieved but should be possible if sufficient resources and time can be allocated for researching this, for which the first step should be to determine the autotrophic partner species of the fungi upon which the *Gymnosiphon* depends. Without this information, planning for the translocation of any achlorophyllous mycotroph has a low chance of success.

###### HISTORY OF THE DISCOVERY

The first record of *Gymnosiphon fonensis* was made by BT in the Boyboyba forest 24 Aug. 2008, but the specimens were erroneously identified as *Gymnosiphon bekensis* by MC. Shortly afterwards at the same site, XVDB discovered two other *Gymnosiphon* species to be present one of which was described as *G. samouritoreanus* (Cheek & van der Burgt 2010) and assessed as Endangered (Couch & Cheek 2019). Subsequently, targeted searches for *Gymnosiphon* to determine the full range of the threatened species found at Simandou led by XVDB later in 2008 found *Gymnosiphon samouritoreanus* (and also *G. longistylus*) at several other sites at Simandou, and also at several sites at Mount Ziama, to the West. However, in contrast, *Gymnosiphon* “bekensis” (here named *G. fonensis*) was not found elsewhere apart from at three other sites at Pic de Fon, Simandou, despite being much larger and more conspicuous in flower than any other West African *Gymnosiphon* species. *Gymnosiphon fonensis* was not recorded again until early Sept. 2022, mainly because no additional surveys were carried out at southern Simandou 2008-2020, and the 2021 project “biodiversity refresh” survey was outside the flowering season of *Gymnosiphon* and of most other fully mycotrophic species. The most recent record of *G. fonensis* was made at a new site 17 Sept. 2022 by AT, when surveying the route for the expected rail route north of southern Simandou.

###### NOTES ON THE BOYBOYBA FOREST

The type location of *Gymnosiphon fonensis* is a small plot c. 30 m x 10 m within the Boyboyba Forest, a small (c. 0.4 km^2^), ecologically unique submontane forest in the Loma-Man Mts, fed by numerous springs, seepages and streams, and which has several other highly threatened species, notably the largest global population recorded for *Keetia futa* Cheek (Rubiaceae), a forest liana globally unique to Pic de Fon, also Critically Endangered (Canteiro & Cheek 2019) and with a nearly identical extant geographical range to that of *Gymnosiphon fonensis*. The Boyboyba Forest also holds the largest global population of an unusual Endangered range-restricted Olacaceae liana, *Anacolosa deniseae* Cheek ined. (Cheek *et al*. 2022b) and contains several other threatened species of plant, and is expected to yield additional new species to science.

The Boyboyba forest has the highest species diversity recorded in West Africa for achlorophllyous mycotrophic species with five species (Cheek & van der Burgt 2010). The five species are: *Campylosiphon* (formerly *Burmannia) congestus* (C.H.Wright) Maas (Burmanniaceae), *Sebaea oligantha* (Gilg) Schinz (Gentianaceae), *Gymnosiphon longistylus* (Benth.) Hutch., *G. samouritoureanus* and *G. fonensis* (Burmanniaceae. All five species grow together in the single plot in which *Gymnosiphon fonensis* is found. Range-restricted species potentially new to science in several animal groups are also in the course of being researched at Boyboyba

## DISCUSSION

Superficially, *Gymnosiphon fonensis* resembles most closely *Gymnosiphon bekensis* of central Africa, and it is possible that the two share a recent common ancestor. The two differ in several other features apart from those documented in Table 1. In *Gymnosiphon fonensis*:

a. The inflorescence is almost always monochasial (vs usually dichasial in *G. bekensis*).
b. The flowers have a short but distinct pedicel to c. 0.5 mm long (vs fully sessile in *G. bekensis*)
c. The anther connective is absent to inconspicuous (vs conspicuous, about as large as the anther cells in *G. bekensis*)
d. The septal nectar glands are only ¼ the length of the ovary cells (vs ½ the length in *G. bekensis*)
e. The seed surface is reticulate (vs verrucose in *G. bekensis*)

However, it cannot be ruled out that *Gymnosiphon fonensis* may instead be sister to *Gymnosiphon samouritoureanus* with which it is sympatric, despite the disparity in size, colour, and shape and posture of the lateral parts of the outer tepal lobes. Both species share sessile or subsessile flowers. They also agree in the deep insertion of the inner tepal lobes, and in the two cells of the anthers being almost adjacent to each other, and not widely separated by the connective. The more or less horizontal, obtriangular stigmas are also similar, as are the proportion of the length of the (minute) nectar glands in relation to the length of the ovary. A molecular phylogenetic analysis is advisable to determine which species is sister to *Gymnosiphon fonensis*. Thus far such studies have included only three (all from Africa) of the ten species now known to occur in Africa and Madagascar (Cheek 2006; Merckx *et al*. 2006).

In the open flower (as studied in spirit preserved material of *Cheek* 19344) the anthers in *Gymnosiphon fonensis* are unusual in the genus in that they appear to be attached to the style head below the stigmatic lobes (see Fig. 2 G&I), having presumably become detached from their site of development on the inner surface of the perianth tube as the flower develops towards anthesis. Therefore, the style head acts as a secondary pollen presenter, as occurs in several other plant families, e.g. Rubiaceae (Cheek & Dawson 2000; Lin *et al*. 2012). So far as we know, this is the first case of secondary pollen presentation recorded in the Burmanniaceae.

The discovery of *Gymnosiphon fonensis*, due to plans for adjacent research facilities, offers the possibility that the mystery of the pollination and seed dispersal, and other aspects of the biology of this pantropical genus, such as the identify of its mycorrhizal associates and their autotrophic partner species, might at last be resolved. To date concrete observations on these for the genus have been completely absent, and only speculation exists (Cheek *et al*. 2006; Merckx 2008).

## Acknowledgements

The specimens and observations cited in this paper were collected with the support of Simfer S.A. and those collected in 2022 under the framework of collaboration between RBG, Kew and Sylvatrop Consulting, and under the Memorandum of Collaboration between RBG, Kew and Universite Gamal Abdel Nasser-Herbier National de Guinee (MINRESI), i.e. UGANC-HNG. We especially thank former Simfer Environment staff John Merry, Leon Payne, Thomas Williams, and current staff Mohamed Tahlaoui, Harry Nevard, M. Soumaoro and M. Diaby. At Sylvatrop, we thank especially Dr Eric Muller and Sylvain Dufour. The authors thank their colleagues at UGAN-C-HNG, and Dr Xander van der Burgt at RBG, Kew for support.

## Notes

### Competing Interest Statement

The authors have declared no competing interest.

